# Simulation of the omicron variant of SARS-CoV-2 shows broad antibody escape, weakened ACE2 binding, and modest increase in furin binding

**DOI:** 10.1101/2021.12.14.472704

**Authors:** M. Zaki Jawaid, A. Baidya, R. Mahboubi-Ardakani, Richard L. Davis, Daniel L. Cox

## Abstract

The recent emergence of the omicron variant of the SARS-CoV-2 virus with large numbers of mutations has raised concern about a potential new surge in infections. Here we use molecular dynamics to study the biophysics of the interface of the omicron spike protein binding to (i) the ACE2 receptor protein, (ii) antibodies from all known binding regions, and (iii) the furin binding domain. Our simulations suggest that while there is significant reduction of antibody binding strength corresponding to escape, the omicron spike pays a cost in terms of weaker receptor binding. The furin cleavage domain is the same or weaker binding than the alpha variant, suggesting less viral load and disease intensity than the extant delta variant.

## Introduction

The omicron variant of the SARS-CoV-2 virus was first detected publicly in Nov. 2021 [1], and traced back to variants which appeared in mid 2020. Because the variant contains a large number of mutations relative to the original strain, including three relevant regions of the viral surface spike protein (the receptor binding domain (RBD), the furin cleavage domain (FCD), and the n-terminal domain (NTD)), the variant is of great concern. There is preliminary evidence that it has overtaken the predominant delta variant in South Africa where it was first detected [2].

The fitness of a particular variant depends upon several factors. First, strong binding to surface receptors is of critical importance, and the SARS-CoV-2 RBD binds with high affinity to the ACE2 protein on human cells [3]. This contrasts with likely weaker binding of coronaviruses associated with the common cold such as OC43 which binds more weakly to sialic acid groups on the cell [4]. Second, escaping the background antibody (Ab) spectrum can confer relative fitness to the dominant variant. Third, efficient membrane fusion and transmission is apparently strongly regulated by the FCD. It has been shown, for example, that ferrets inoculated with a WT SARS-CoV-2 with the FCD deleted can become infected but fail to transmit to other ferrets [5]. The high viral load of the delta variant has been clearly associated with the mutation P681R of the FCD [6] and has led to the current dominance of 96% of SARS-CoV-2 sequences worldwide [7].

Given the time lag in carrying out protein synthesis, structure determination of bound complexes, determining protein binding affinities, and measuring viral neutralization by Abs for new variants, there is clearly a role for rapid computational studies that can assess the differences of new variants relative to background variants as they arise.

In this Letter, we point out here that computational *ab initio* molecular dynamics studies of omicron RBD-ACE2, RBD-antibody (AB), FCD-Furin, and NTD-antibody are consistent with: 1) robust antibody escape in all regions compared to wild type (WT) and delta, 2) FCD binding to Furin intermediate between WT and delta, and 3) weaker binding to the ACE2 than WT or delta. The Ab escape can confer transmissibility advantages for a population with a prevalent delta variant Ab spectrum, but the weaker binding to ACE2 and modest enhancement of furin binding are likely to lead to weaker transmissibility than delta. This work uses ColabFold’s [8] implementation of AlphaFold-Multimer [9] to generate structures for FCD-furin binding.

## Materials and Methods

### 0.1 Molecular Models

We drew our starting structures for RBD-ACE2 binding from PDB file 7A94 [10] reference for the WT and delta variants, and PDB file 6M0J [11] for the omicron variant. For Class I ABs, which bind in the same region of the RBD as the ACE2, we used 7CJF [12] and 7KVF [13], while as a representative class III Ab that binds to the RBD away from the ACE2 interface, we used 6YOR [14]. For an NTD-Ab we used 7C2L [15].

The antibodies chosen do not comprehensively portray all neutralizing Abs for the SARS-CoV-2 spike, but are representative of the spectrum of antibodies that neutralize the SARS-CoV-2 virus. This study does not account for t-cell binding sites [16]. Fig. 1 shows the structures of the different complexes studied in this paper.

**Figure 1.**
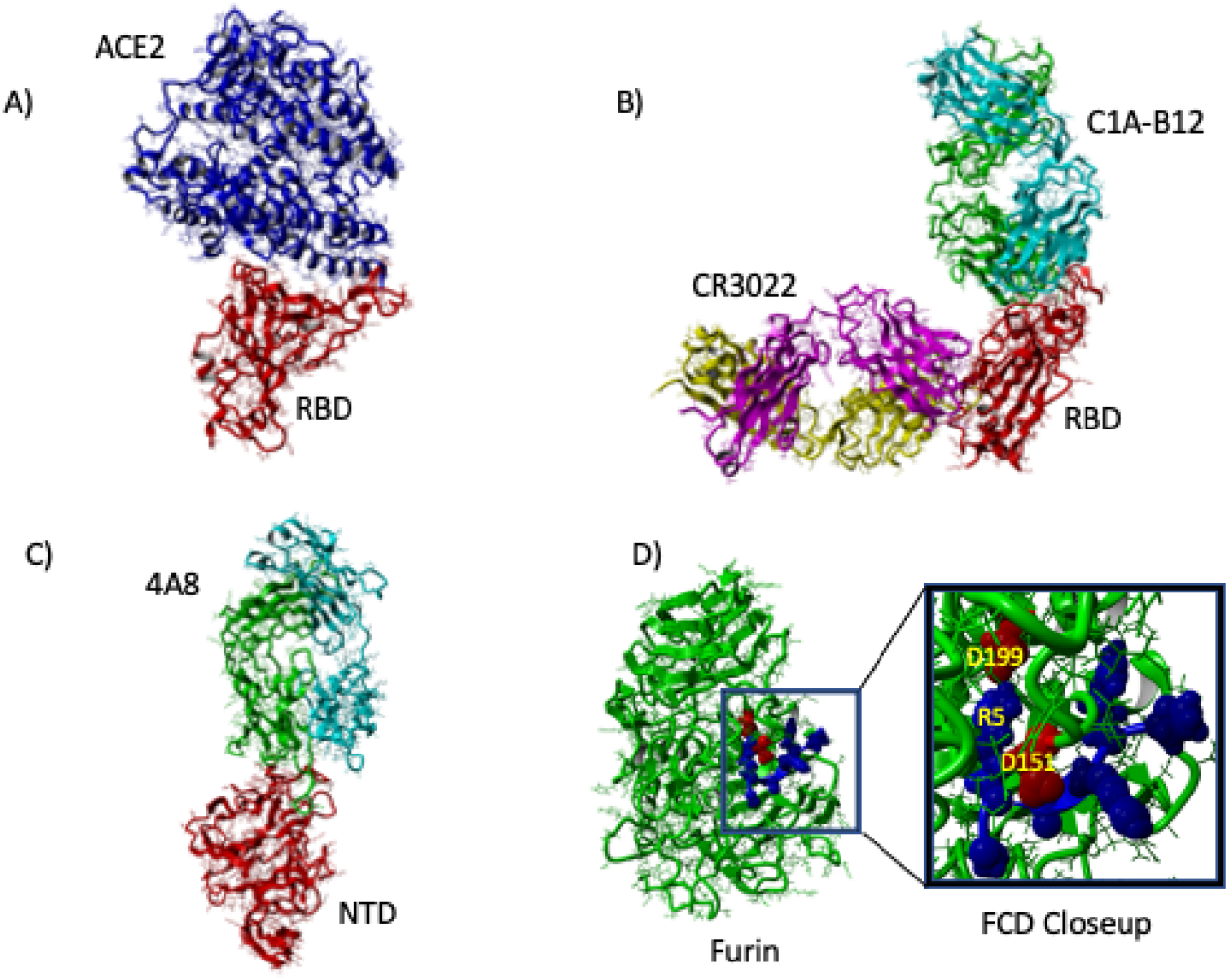
Structures of WT spike protein complexes studied. A) ACE2(red)-RBD(blue) binding. B) Binding of RBD(red) to Class I Ab C1A-B12 (binds in ACE2 interface region; heavy chain green, light chain cyan) and Class III Ab CR3022 (binds away from ACE2; heavy chain magenta, light chain yellow). C) Binding of NTD to 4A8 Ab (heavy chain green, light chain cyan). D) Binding of FCD (blue) to furin (red). Blowup highlighting position of fifth residue R5 (R685 for WT SARS-CoV-2) with proximate aspartic acid residues D151, D199 of the furin enzyme.

### 0.2 Molecular Dynamics

To simulate the protein-protein interactions, we used the molecular-modelling package `YASARA` [17] to substitute individual residues and to search for minimum-energy conformations on the resulting modified structures of the complexes listed in Table 1. For all of the structures, we carried out an energy-minimization (EM) routine, which includes steepest descent and simulated annealing (until free energy stabilizes to within 50 J/mol) minimization to remove clashes. All molecular-dynamics simulations were run using the AMBER14 force field with [18] for solute, GAFF2 [19] and AM1BCC [20] for ligands, and TIP3P for water. The cutoff was 8 Å for Van der Waals forces (AMBER’s default value [21]) and no cutoff was applied for electrostatic forces (using the Particle Mesh Ewald algorithm [22]). The equations of motion were integrated with a multiple timestep of 1.25 fs for bonded interactions and 2.5 fs for non-bonded interactions at *T* = 298 K and *P* = 1 atm (NPT ensemble) via algorithms described in [23]. Prior to counting the hydrogen bonds and calculating the free energy, we carry out several pre-processing steps on the structure including an optimization of the hydrogen-bonding network [24] to increase the solute stability and a *pK*_a_ prediction to fine-tune the protonation states of protein residues at the chosen pH of 7.4 [23]. Insertions and mutations were carried out using `YASARA`’s BuildLoop and SwapRes commands [23] respectively. Simulation data was collected every 100ps after 1-2ns of equilibration time, as determined by the solute root mean square deviations (RMSDs) from the starting structure.

**Table 1.** Interfacial hydrogen bonds between proteins for WT, delta, and omicron. Last column: reference PDB structure

| Bound Pair | WT | delta | omicron | PDB |
| --- | --- | --- | --- | --- |
| RBD-ACE2 | 8.43±1.69 | 8.32±1.46 | 5.06±1.20 | 7A94(WT,delta)<br>6M0J |
| RBD-P4A1<br>(Class I) | 17.76±2.0 | 17.61±2.03 | 13.9±1.89 | 7CJF |
| RBD-C1A-B12<br>(Class I) | 19.23±1.87 | 14.94±2.66 | 11.41±2.49 | 7KFV |
| RBD-CR3022<br>(Class III) | 12.59±1.82 | 12.0±1.53 | 10.85±1.43 | 6YOR |
| NTD-4A8 | 9.62±1.78 | 7.48±1.49 | 4.73±1.63 | 7C2L |
| FCD-Furin | 10.93±1.72 | 15.36±1.86 | 8.65±1.59 (K679)<br>12.05±1.68 | NA |

The hydrogen bond (HBond) counts were tabulated using a distance and angle approximation between donor and acceptor atoms as described in [24]. Note that in this approach, salt bridges of proximate residues, are effectively counted as H-bonds between basic side chain amide groups and acidic side chain carboxyl groups.

### 0.3 Endpoint Free Energy Analysis

We calculated binding free energy for the energy-minimized structure using the molecular mechanics/generalized Born surface area (MM/GBSA) method [25–27], which is implemented by the HawkDock server [28]. While the MM/GBSA approximations overestimate the magnitude of binding free energy relative to *in-vitro* methods, the obtained values correlate well with H-bond counts. For each RBD-ACE2, RBD-AB, and NTD-Ab binding pair we average over five snapshots of equilibrium conformations. For each FCD-furin pair, we average over ten snapshots of equilibrium conformations.

### 0.4 Use of ColabFold/AlphaFold for Furin binding domain

We will present a complete description of the use of ColabFold/AlphaFold for modeling the FCD-furin binding separately [29]. Full details of this method are provided in [8, 9, 30]. In brief, we used the heterocomplex prediction method known as AlphaFold-Multimer [9, 30] as implemented within ColabFold [8] to predict the best bound structure to the furin enzyme of the six residue FCD from the WT protein. The delta and omicron structures were then obtained by mutation from the predicted WT FCD-furin structure.

## 1 Results

### 1.1 Binding Strengths: HBond and Binding Free Energy

Our main results for interfacial HBonds are summarized in Table 1. We find weaker binding to the ACE2 receptor in contrast to a recent quantum calculation [31], which should moderate infectivity, and significant antibody escape of the omicron for all three regions (Class I, Class III, and NTD) considered. This escape is measured by the reduction in hydrogen bond count between the antibodies and the spike protein. For the FCD-Furin binding, six residues fit into the binding pocket, which we argue elsewhere to begin with residue 681 for WT, alpha, and delta. For omicron, we consider the possibility of leading with the N679K mutation or P681H mutation. The latter is the same as the alpha variant. We see that the expected binding to the FCD is at best the same as the alpha variant, and significantly less than the delta variant.

The binding energies from the GBSA analysis of molecular dynamics equilibrium conformations are shown in Table 2. The same PDBs are utilized. Evidently the trend of binding energies tracks well with the easier to estimate interfacial HBond count, with the lone exception of the Class III Ab binding to the delta variant.

**Table 2.** GBSA Binding free energy estimate in kcal/mole

| Bound Pair | WT | delta | omicron |
| --- | --- | --- | --- |
| RBD-ACE2 | -62.9±6.8 | -64.3±3.6 | -55.5±4.5 |
| RBD-P4A1<br>(Class I) | -115.9±7.0 | -110.2±7.5 | -75.6±9.5 |
| RBD-C1A-B12<br>(Class I) | -77.9±8.0 | -70.3±8.4 | -50.8±5.6 |
| RBD-CR3022<br>(Class III) | -95.4±2.5 | -111.7±7.8 | -84.1±13.3 |
| NTD-4A8 | -88.9±17.9 | -81.1±5.7 |  |
| FCD-Furin | -83.6±8.4 | -117.3±4.8 | -63.5±3.6 (K679)<br>-93.2±3.8 |

### 1.2 Mutations leading to Ab Escape and weaker ACE2 binding

Fig. 2 illustrates the key mutations leading to differences in binding for the delta and omicron variants relative to WT.

**Figure 2.**
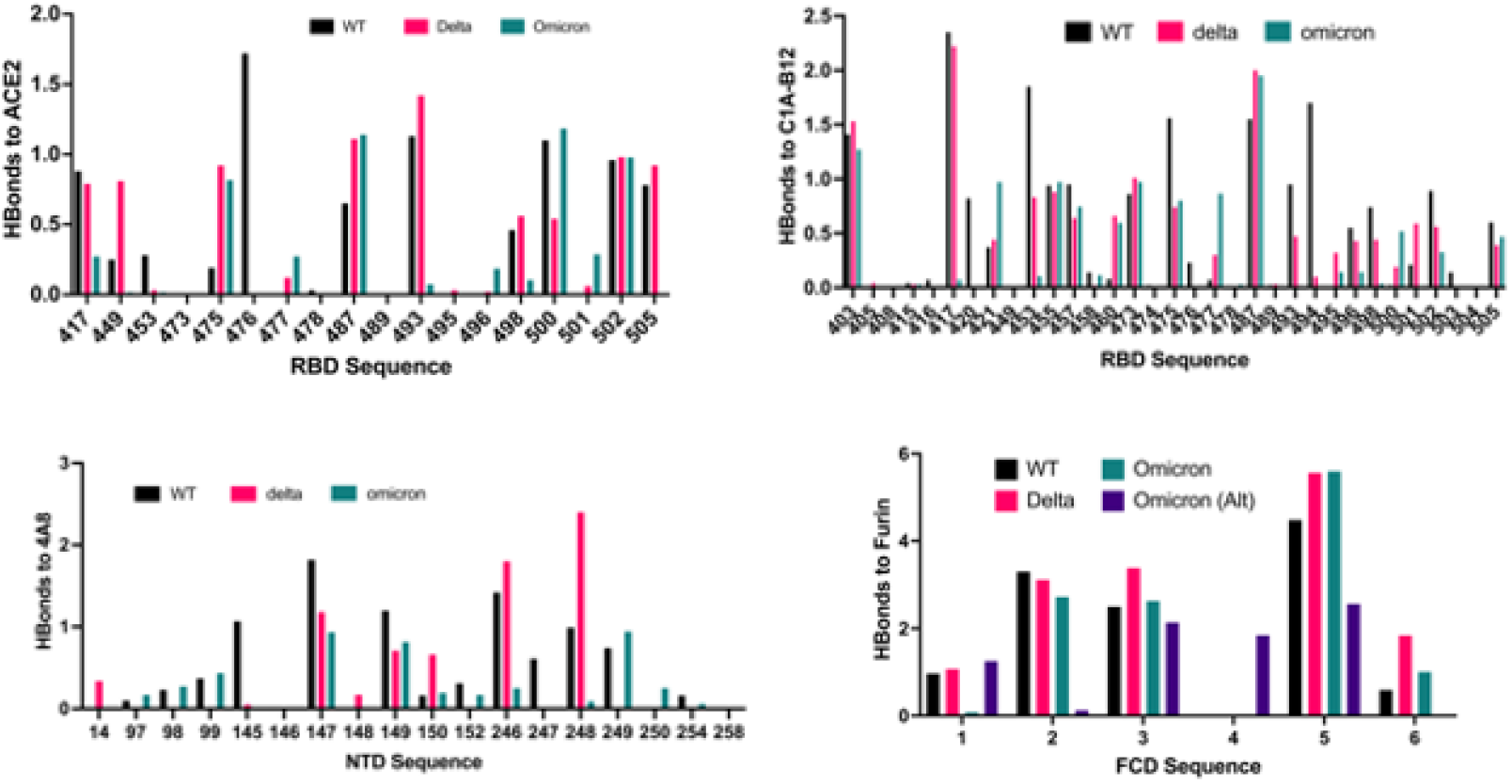
Overview of binding changes for delta and omicron variants relative to WT. Color coding is the same for all charts. For the FCD to furin binding, R1-R6 correspond to 681-686, except for the alternate omicron sequence 679-684. For clarity, RBD binding to P4A1 and CR3022 Abs are not shown.

*ACE2* For ACE2 binding, four key mutations weaken the ACE2 binding for omicron: 1) K417N removes the K417(RBD)-D30(ACE2) salt bridge. 2) Q493K removes hydrogen bonding between the glutamine side chain and residues K31 and E35 on the ACE2 driven by K-K coulomb repulsion. 3) Q498R removes hydrogen bonding between the glutamine side chain and K353 of the ACE2 driven by R-K Coulomb repulsion. 4) Y505H removes hydrogen bonding between the Y505 sidechain where the O acts as a donor and the E37 sidechain of the ACE2.

*Class I Abs* For Class I antibodies, the following mutations are critical to reducing binding strength: For binding to P4A1, 1) the Y455 binding to Y33.HC of the Ab heavy chain (HC) is removed. 2) The Q493K, G496S, and Q498R mutations lead to removal of bonds with E101.HC, W32.LC of the Ab light chain, and S67.LC. 3) The Y505H mutation removes bonds to S93.LC. For binding to C1A-B12, 1) the K417N mutation removes a salt bridge to D96.HC, a side chain bond to S98.HC, and weakens a side chain bond to Y52.HC. 2) The mutations Q493K, G496S, and Q498R remove bonds to R100.HC, S30.HC, and S67.HC. 3) The N501Y and Y505H mutations weaken bonds in the 501-505 region to G28.LC, S30.LC, and S93.LC.

*Class III Ab* For the Class III antibody CR3022, the most noticeable differences compared to WT are 1) the absence of binding at N370 to Y27.HC. This appears to be driven by the hydrophobic substitution S371L, which pulls the asparagine at 370 out of bonding distance from Y27.HC. 2) Weakened bonding of T385 to S100.HC.

*NTD Ab* For the NTD Ab 4A8, we find that the notable differences of omicron compared to WT are 1) weakened binding at 145-152 presumably due to the deletion at 142-145 relative to WT, and 2) significantly weakened bonding at 246-254 driven by the EPE insertion at 214 and the deletion at 211. Both the 142-145 deletion and the 211 deletion with EPE insertion disrupt the epitope positionings at 145-152 and 246-254 respectively.

### 1.3 Mutations in the FCD

As we will discuss in more detail elsewhere, for the generic 681-686 sequence of the FCD, the most critical residue apears to be the 685. In the WT, the arginine is able to form a salt bridge in the interior pocket with D199 of the furin, and bond additionally with S146, W147, D151, A185, and S261. This tendency is illustrated in Fig. 1D. These bonds are all strengthened for delta and omicron. For the alternate KSHRRA sequence of the omicron, beginning at 679, the position of the arginine in the binding pocket allows only the salt bridge formation with D199.

As shown in Fig. 3, we observe that the binding strength, which is determined to a large degree by the binding of the fifth residue of the FCD, correlates inversely with the root mean square fluctuation (RMSF) of the backbone C*α* of the first FCD residue at 681. This suggests that locking the 681 C*α* as happens for P681R is a key to lowering the fluctuation spectrum of the 685 residue allowing for stronger binding at this site. Evidently, the gain in binding enthalpy offsets any advantages in conformational entropy for the FCD.

**Figure 3.**
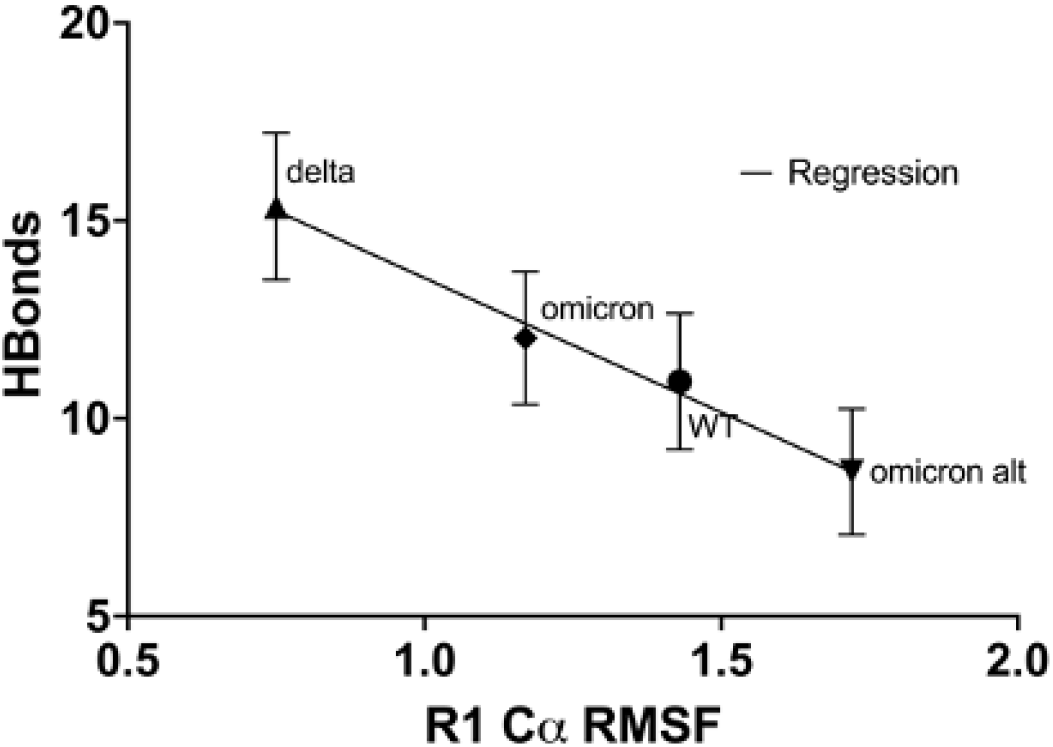
Correlation of FCD-furin interfacial HBond count with RMSF of first residue in FCD. The higher the RMSF of the first residue in the FCD, the harder it is to bind to the furin, especially for the critical fifth residue which inserts into the furin pocket as shown in Fig. 1D. R1 is residue 681 for all but the alternate omicron sequence which starts at residue 679.

## 2 Discussion

We observe that the strength of binding of the FCD appears to correlate well with the fluctuations of the initial residue at 681. The lower the fluctuation of the backbone carbon, the lower the fluctuation of the backbone carbon for residue 685, which dominates the bonding to the furin. The P681R mutation provides the lowest C*α* RMSF observed amongst the four FCD examples considered here, and the alternate K679 starting point for omicron provides the largest C*α* RMSF.

The lower severity of omicron versus delta is related to the Furin Cleavage Domain. Once a corona virus is bound to a human cell, disease severity is regulated by furin cleavage of the spike. After initial binding to the human ACE2 protein, Furin protease cleavage breaks the spike to facilitate cell wall fusion [6] and viral reproduction. The higher the furin-FCD HBond binding count, the more efficient the fusion at the molecular level, and ultimately, higher viral load on the host.

In summary, a consistent picture of omicron in comparison to the delta strain is emerging. Hospitalization data points to higher disease transmissibility but lower severity for the omicron strain compared to delta [32]. Our simulations see lower interfacial HBond counts for omicron for known RBD and NTD binding regions consistent with this, as well as weaker ACE2 binding and furin binding than the delta variant. Against an immunity background tuned to the delta variant, omicron will be more transmissible. (Note that two other studies suggest stronger ACE2 binding from omicron [31, 33].) Experimental studies of the binding of the RBD to ACE2 and the FCD to furin will be needed to confirm these predictions.

## Acknowledgments

We acknowledge useful conversations with Javier Arsuaga, Victor Muñoz, and Mariel Vazquez.

